# A spatially targeted metric of potential koala populations based on Bayesian Networks for conservation management

**DOI:** 10.1101/2025.05.20.655061

**Authors:** Grace Muriuki, Shannon Mooney, Michael Castiglione, Jac Davis, Jegar Pitchforth, Sandra Johnson, Kalpani Ishara Duwalage, Kerrie Mengersen, Paul Pao-Yen Wu

## Abstract

Koalas (Phascolarctos cinereus) face significant threats from habitat loss and fragmentation, with South**-**East Queensland (SEQ) experiencing some of the most severe population declines. Current metrics used to assess conservation impacts often lack spatial resolution, transparency, or integration of diverse data sources. We propose a novel, spatially targeted metric, Koala Population Potential (KPP), to estimate the potential number of koalas that could inhabit a given area under ideal conditions. This metric is derived from a Bayesian Network model that integrates ecological, environmental, and threat data, including satellite imagery, expert knowledge, and stakeholder input. We developed and validated the KPP within an interactive online platform, the Koala Bayesian Network Platform (KBNP), enabling stakeholders to visualise and assess the conservation impact of various management scenarios at the land-parcel scale. Application of the KPP across SEQ revealed substantial spatial variation in potential koala populations. Regional areas like Somerset and the Scenic Rim exhibited high KPP scores, while urban LGAs such as Redland City and Logan had lower scores due to elevated threat levels. Model predictions of habitat support were strongly correlated with independent habitat quality assessments (R² = 0.803, p < .001), supporting the validity of the approach. The KBNP provides an accessible, evidence-based decision-support tool that empowers landholders, planners, and conservation managers to evaluate and communicate the likely outcomes of their actions. While developed for koalas, this framework is adaptable to other vulnerable species requiring localised, data-driven conservation planning under uncertainty.

## Introduction

Fifty years ago, the Australian koala (*Phascolarctos cinereus*) was abundant across Australia. In 2012, the Australian Government estimated that populations in the State of Queensland were declining at a rate of 53 percent each koala generation (6-8 years). In February 2022, the koala was listed as Endangered in Queensland [1]. Habitat loss, degradation, and fragmentation remain among the most significant threats to koala populations, particularly in rapidly developing areas like South-East Queensland (SEQ). These pressures, compounded by other threats such as climate change and disease, have been identified by both researchers and local authorities as key drivers of koala population decline [2], [3]. This is a stark example of a global phenomenon of species decline and underscores the urgent need for effective habitat support and innovative conservation approaches. To better address these challenges, new methods for continuous monitoring, rapid impact assessment, and real-time tracking of conservation actions are critical.

One of the major challenges facing conservation is the limited ability of established systems to accurately measure and track the target species [4], [5]. Stakeholders - including natural resource managers, developers, and government - face the lack of an accepted metric for estimating the impacts of development or conservation actions, as well as the resultant impacts of these developments on populations and long-term viability of the remnant populations. In this regard, a metric and underpinning model that is able to link actions and their impacts to the factors necessary for sustaining Koalas populations is key to supporting management actions for conservation. A population potential metric provides a practical alternative by estimating how many koalas could be supported in an area under ideal conditions, allowing managers to track habitat suitability and viability over time even when direct population data are scarce or unreliable.

A second challenge is that data that can provide an evidence base for monitoring and impact assessment are often limited and disparate. For example, koalas are typically elusive and difficult to observe directly [6], [7], and active collection of wildlife sighting data by trained experts is expensive and prone to false negatives where a species is not detected when it is indeed present [5]. Other approaches focus on directly measuring koala survival, through tracking individuals or through veterinary records from wildlife hospitals but these approaches also have limitations [6], [8], [9]. There is global interest in the use of other sources of information such as satellite imagery and citizen science for species counts and geolocation, but these data may be inaccessible for many stakeholders. While availability of data in the requisite spatial and temporal scales remains a challenge, integration of currently available data in a format that also allows ongoing improvements and opens opportunity for integration as better data for Koala conservation becomes available. To address these challenges, a population potential metric, utilising advanced statistical models like Bayesian Networks, can integrate diverse data sources, providing a dynamic and adaptable framework that evolves as new information is incorporated.

A further challenge is the lack of accessible, explainable, publicly available digital platforms for interactive evaluation of costs, benefits and consequences of potential conservation actions [10]. These limitations mean that for many stakeholders it is not practical to evaluate the impacts of their actions on species numbers, and that opportunities for comparing information, sharing insights, and collaborative learning are limited. An enduring challenge in conservation is the lag between research and its application by practitioners; for a threatened species, the urgency necessitates that those implementing actions for its conservation have access to tools to make decisions quickly, and to contribute to the improvement of these tools. Embedding a population potential metric into a user-friendly platform allows stakeholders to rapidly assess and visualise the outcomes of their conservation efforts, even when underlying data are uncertain.

Conventional approaches like current population estimates and carrying capacity have been widely used in koala conservation, but they pose several limitations. Population estimates often rely on outdated or incomplete data, limiting their accuracy and relevance [11]. Traditional carrying capacity models assume fixed ecological limits and often overlook important changes in the environment over time, such as seasonal variation, land-use change, or climate effects, which can significantly influence species distribution and habitat quality [12]. Additionally, both methods often lack the spatial granularity needed for local decision-making [13]. These limitations emphasise the need for a more dynamic, scalable, and spatially explicit metric, such as the Koala Population Potential proposed in this study.

In this paper, we propose a new metric called Koala Population Potential (KPP) for describing and evaluating the impact of monitoring, impact assessment and conservation actions for vulnerable species. This metric is based on an underlying Bayesian Network (BN) that captures the complex system of factors affecting KPP, fusing modern data sources, including satellite data, surveys, citizen science data and stakeholder knowledge, to produce a ‘population potential’ for the target species. This indicates the possible number of the target species that could potentially thrive in a given area under specified conditions. The metric is delivered through an innovative user-focused online Koala Bayesian Network Platform (KBNP). Interactive toggles embedded in the platform enable managers, researchers and land-holders to evaluate the impact of their own actions on the species population potential at a scale appropriate for decision-making. Here, the scale adopted is the land parcel size, since this indicates carrying capacity and is employed by different decision makers including local governments, businesses and landholders. We describe and demonstrate our approach in the context of the urgent need for a rapid, remote impact assessment system for koala conservation in SEQ. While the model proposed in this paper does not directly measure habitat loss, it incorporates several indirect indicators, such as land use type and spatial context, that reflect habitat condition and its influence on population viability.

The paper proceeds as follows. The new species population potential metric (KPP) is presented in the Materials and Methods section, in the context of koala conservation in SEQ. The application of the metric under a range of conditions and for a variety of real-world scenarios is described in the Results section. The paper concludes with a general Discussion.

## Materials and Methods

### Overview of the koala study

To date, expert elicitation has been used as a measure of koala population due to the extremely challenging nature of making observations of koalas. In 2012, an expert elicitation estimated that the national koala population size was 329,000 (range: 144,000-605,000) [14] - In 2012, this expert elicitation estimated that that there were 79,264 koalas in Queensland and SEQ (15,821). Based on 2012 population estimates [14], the bioregions with the highest density of koalas in Queensland included SEQ (SEQ, 0.002 koalas/ha).

The current project builds on previous work assessing habitat suitability for koalas across SEQ. The Queensland Department of Environment and Science (DES) released a South-East Queensland Koala Conservation Strategy [15] that included spatial modelling identifying areas of existing koala habitat, locally important vegetation, and habitat restoration areas. A DES - funded habitat assessment toolkit used satellite data to provide habitat metrics for public and private landholders in Queensland [16]. The goals were to stabilise koala populations in SEQ, a net gain in core koala habitat area, rehabilitation to habitat, and a 25 per cent reduction in disease, injury, and mortality [15].

### A solutions-focused outcome metric: KPP

Although the concept of koala survival is simple in principle, in practice there is no strong consensus on how to measure koala populations in the wild. Various definitions are provided in Table 1.

**Table 1.** Current operational definitions for number of koalas.

| Indicator | Definitions |
| --- | --- |
| Observed number of koalas | Wildlife hospital records [6], [17]<br>Thermal imaging [7]<br>Tracked survival rates [8]<br>Citizen science reports [5], [18] |
|  | Koala surveys [19]<br>Acoustic sensors [20] |
| Estimated number of koalas | Expert-estimated number of future koalas [21]<br>Management effect on future koala numbers [22] |
| Habitat biophysical traits | Proportion of primary Eucalyptus tree species [12]<br>Soil fertility [21], [23] |
| Carrying capacity | Number of animals the landscape can support [24] |

An alternative solution is to focus on expert estimations of current and future potential numbers of koalas under different management scenarios [21]. We drew on existing concepts of koala survival, and practical considerations relating to time and cost of constructing a koala management decision support system. Also considered was the purpose of the decision support tool: while decision-makers may not be able to influence the actual number of koalas in an area, they can directly influence the conditions to be either friendly or hostile to koala populations. Hence, we defined koala survival as “*the potential number of koalas that could exist in an area, if all conditions were ideal*”.

Our operationalisation of this definition builds on prior research into habitat suitability, threat modelling, and carrying capacity [12], [21], [23], [24]. We model KPP using three spatially explicit variables that are broadly supported in the literature: the land area available (A), the quality of the habitat (H), and the level of threats (T). These variables capture the landscape’s potential to support koalas, accounting for both natural habitat conditions and human-induced threats.

The resulting KPP score represents the number of koalas that could potentially inhabit a given area under ideal conditions, constrained by habitat quality and threat levels. There is uncertainty around this estimate, based on the amount and quality of available data. This uncertainty varies across regions and is captured through a credible interval which indicates that, with a given degree of confidence, the true level of KPP lies within this range, given the data. For many applications, instead of focusing on the aggregate KPP score, it may be more useful to focus on the credible interval containing the range of possible KPP values for each region. In addition to providing intrinsic information about an area, these indicators can also be used to make comparisons between geographic areas and track koala conservation efforts over time.

Formally, the KPP score is defined as ‘the largest possible number of koalas in a given area, modified by limitations of habitat and threats’.

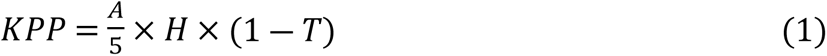

where:

A = The land area in hectares
H = The level of habitat support (0,1) within the land area where 1 is a perfect koala habitat, and 0 is a habitat that provides no support to koalas (e.g. ocean).
T = The level of threats in the land area (0,1) where 1 is the highest possible level of threat, and 0 is no presence of threats.

Experts have indicated that this definition of KPP is aligned with the existing thinking around koala management, and that the assumption of a maximum of 1 koala per 5 hectares is supported by the current literature (See Results section).

The KPP differs from existing metrics in the following ways:

#### Carrying capacity

KPP is similar to the ecological metric of “carrying capacity” [24] but is intended as a more flexible term with a wider definition. Carrying capacity is reflected by the first element (A/5*H) of KPP, which is KPP with modification based on habitat condition, but without correction for the presence of threats. As such, the KPP of a given area should always be equal to or lower than its ‘carrying capacity’.

#### Species presence models

KPP also stands in contrast with species presence models aimed at estimating the true number of koalas. Species presence models, especially for a cryptic species such as the koala, require a substantial time and financial commitment. Additionally, we did not wish to duplicate the efforts of ongoing projects aiming to estimate extant koalas, for example, the National Koala Monitoring Program [11]. As such, presence modelling outputs should, on average, be distributions centred around the KPP of the same area.

#### Cost-based metrics

KPP does not include any consideration of the cost of implementing a management action. By focusing on outcomes, rather than costs, KPP encourages users to draw upon the specific strengths in their organisations and does not assume that the same actions will have the same costs for every user. KPP can therefore be used as a planning tool for stakeholders to assess the impact of using whatever skills and resources are available to them, empowering users to add that information to their decisions independently.

The Population Potential metric was developed through a BN model. A BN is a type of complex systems model that comprises a set of nodes (representing indicators) and directed arrows (representing connections between the indicators), leading to a desired outcome [25]. We elected to follow a common approach of quantifying each node in the BN through a conditional probability table (CPT) that describes the probability of a discrete outcome (True/False, High/Medium/Low, etc.) based on the values of the ‘parent’ nodes (those indicators that directly influence the particular node through the directed arrows). Such a discrete approach is useful in contexts such as koala conservation with limited data and resolution of data, where categories better reflect the precision of the available data. Additionally, discrete states can be chosen to reflect ecological thresholds or thresholds for management action, allowing for explicit modelling of conservation outcomes or actions of interest [26].

Stakeholders were deeply engaged throughout the project. The goals of this engagement were to elicit a broad overview of user attitudes to koala conservation and understand current workflows and gaps in knowledge, frame user stories for the BN, and evaluate and validate the developed KBNP by generating estimates of the important variables for koala conversation. Engagement activities included online surveys, virtual workshops with topic experts, and in-person workshops with koala conservation practitioners.

The workflow underpinning the creation of the BN followed five steps: (1) Identify the major indicators affecting koala vulnerability; (2) Develop the structure of the BN by creating a systems diagram that depicts connections between the indicators; (3) Quantify the probability tables in the BN using the various data sources; (4) Run the BN to assign baseline koala scores for each land parcel in the area; (5) Validate the model. These steps are depicted in Fig 1 and described in more detail below.

**Fig 1.**
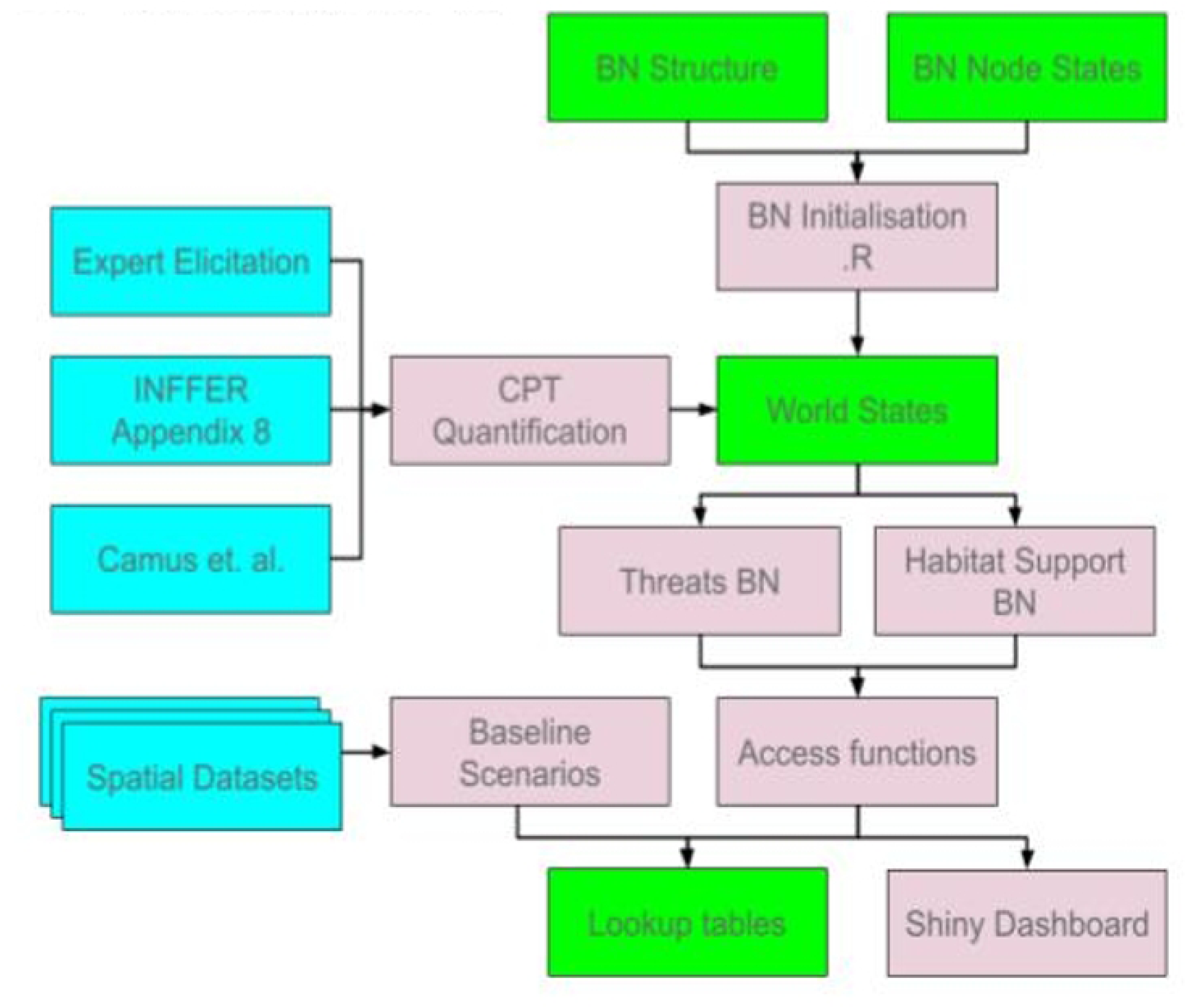
Workflow for constructing the BN model.

##### (1) Major indicators

We relied on previous research to identify indicators of koala survival. Data for these indicators, or a reasonable proxy, was required to be available at a spatial scale and able to be aligned with the unit of analysis for the project. Building on this literature, specific indicators to include in the systems model was informed by previous work on koala habitat suitability conducted by Healthy Land and Water (HLW) [27] and previous work on vulnerability to threats conducted in New South Wales [21], with the guidance and participation of three professionals with technical expertise in koala conservation and GIS methods.

##### (2) Structure of the BN

Based on results of stage (1), a systems diagram depicting the connections between the koala conservation indicators was constructed as a foundational BN model. The network was refined to follow rules for model parsimony [28]: 4 or fewer levels in the model; and 3 or fewer parent nodes for each child node. The diagram was shared with the experts and technical team for feedback and also tested conceptually using two example scenarios: fire and habitat connectivity. The final BN structure was created in light of these results.

##### (3) Quantification of the BN

Satellite imagery was used to establish baseline measurements of relevant environmental indicators for each geographic region. These reference baselines served as a counterfactual or “business as usual” scenario, against which the impact of various scenarios could be estimated [6]. Baseline data were parameterised according to methods established in the Biodiversity Assessment and Mapping Methodology [29]. GIS raster layers representing each variable in the model were processed using the R library sf. Layers were discretised to match the network parameters, as described in Table 2 (see Results). Nodes were restricted to two parameter levels, “low” and “high”, for ease of implementation in the user-facing software.

**Table 2.** Node names, definitions, states, and data.

| Node | Description | States | Data |
| --- | --- | --- | --- |
| Koala Potential (KP) | The number of koalas that could be potentially supported after accounting for habitat support and threat impact | Calculated | $PP * HS * (1-TL)$ |
| Habitat Support (HS) | The suitability of the habitat before threats are considered | Range 0-1<br>1 = highly suitable | Expert Elicitation |
| Parcel Potential (PP) | Theoretical max population capacity of the parcel if all conditions were perfect for koalas | Calculated | 5 koalas per ha |
| Threat Level (TL) | The overall risk level of all threats to the Koala population | Range 0-1<br>1 = high threat level | Adapted from [21] |
| Habitat Size | EHP (Department of Environment and Heritage Protection) category: size of tract (discrete area) of habitat in good condition | Low, High | HLW Vegetation Indices |
| Habitat Connectivity | EHP category: Strength of vegetation clustering on the land package | Low, High | HLW Vegetation Indices |
| Ecology | Probability that the ecology of the land package is suitable for Koalas | Low, High | Latent variable |
| Vegetation Cover | Presence of regrowth or remnant vegetation | Low, High | MCOVER Remnant and Regrowth |
| Dogs | Risk of attacks by domestic dogs | Low, High | QLD Gov. urban footprint |
| Disease | Risk of disease | Low, High | Koala Hospital records |
| Vehicle | Risk of vehicle strike | Low, High | Main Roads wildlife incidents |
| Fire | Risk of fire | Low, High | Wildfires_SEQ_2019 |

Each continuous indicator in the BN was discretised into classes. Summaries for each spatial indicator were created along with suggestions for cutoff values. Two expert elicitation sessions (7 November 2022 and 16 January 2023) were held in which experts ranked the selected parameters from “most important” to “least important” for koala populations. We combined these rankings linearly. Missing combinations of CPTs were estimated based on a computer inference algorithm written by author JP.

##### (4) KPP score

Point estimates for predicted habitat support, threat level, and potential koalas, were calculated for each land parcel in the area using an exact method created by author PW in R statistical software. To calculate the uncertainty of this estimate, habitat support, and threat level were predicted 20 times for each land parcel, conditional on the baseline data, using the cpquery function in the R package bn_learn.

##### (5) Model validation

Face validity for the network model was established via short questionnaire aimed at experts and the project team, administered in January 2023. Convergent validity [30] was established via visual comparison with source material BN for koala survival in NSW [21] and koala habitat suitability models provided by HLW. To support these visual tests, predictive validity was tested by comparing our results to an independent set of scores created by HLW with support from DES to measure a land parcel’s suitability for koala habitat. We compared our habitat support predictions to the corresponding score for each land parcel in SEQ.

### An interpretable, interactive BN Platform: the KBNP

The primary purpose of translating the Koala Bayesian Network to a technology platform is to allow learning between agencies and from experience. In addition to the statistical model, we produce a visible, interactive response to justify the resources expended on the model’s creation and to encourage stakeholders to use it and contribute data. The most important function of a statistical model is to use the results of data analysis to create and disseminate recommendations for change.

Drawing from previous work and consortium experience, and in collaboration with scientific experts and stakeholders, the KBNP consortium worked together to advance the development of a technology platform to allow stakeholders to interact with the Koala Bayesian Network model. The goal of the KBNP is to be used as a shared learning platform by conservation institutions so that the planning, review, comparison, and analysis of actions affecting koalas can be performed rapidly and without specialist knowledge, while retaining confidence in the scientific validity of the methods and data. This technology is also a useful template for new scientific model platforms applied to other flora and fauna conservation projects with insufficient existing data and assessment methods.

## Results

### KBNP development

Relevant indicators for inclusion in the KBNP were drawn from published indicators related to koala survival (see Table S1 in Supporting information), as well as indicators identified by HLW as part of their multi-criteria analysis process, Investment Framework for Environmental Resources (INFFER) is a tool designed to help prioritize and assess environmental projects based on their cost-effectiveness, feasibility, and expected outcomes, and the habitat metrics for public and private landholders in Queensland identified as part of the DES-funded habitat assessment toolkit for koalas [16]. The final list of indicators is presented in Table 2, along with definitions and the states used in the model.

The systems diagram that shows the integration of these indicators in the BN model is depicted in Fig 2.

**Fig 2.**
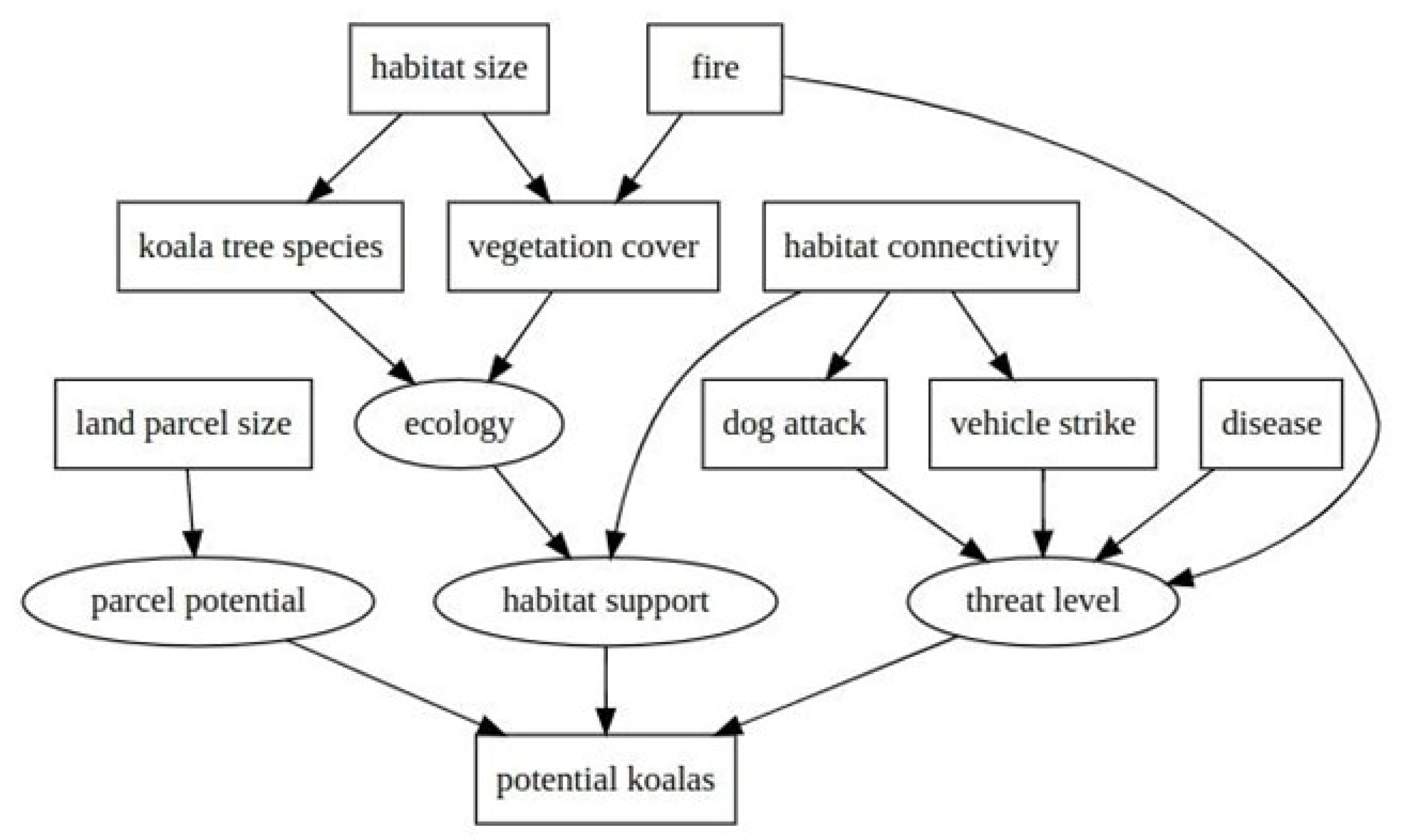
Structure of the Koala Bayesian Network model structure underpinning the new metric, KPP.

Data were derived from expert elicitation for conditional probability distributions and using satellite imagery for georeferenced baseline estimates. The final BN model addressed two main subnetworks: habitat quality (including habitat size, habitat connectivity, vegetation cover, and presence of koala-supporting tree species) and threat level (including presence of dogs, vehicle strikes, fire, and disease). The components of the KBNP that enable visualisation of the BN and interactive engagement with the model is shown in Fig 3.

**Fig 3.**
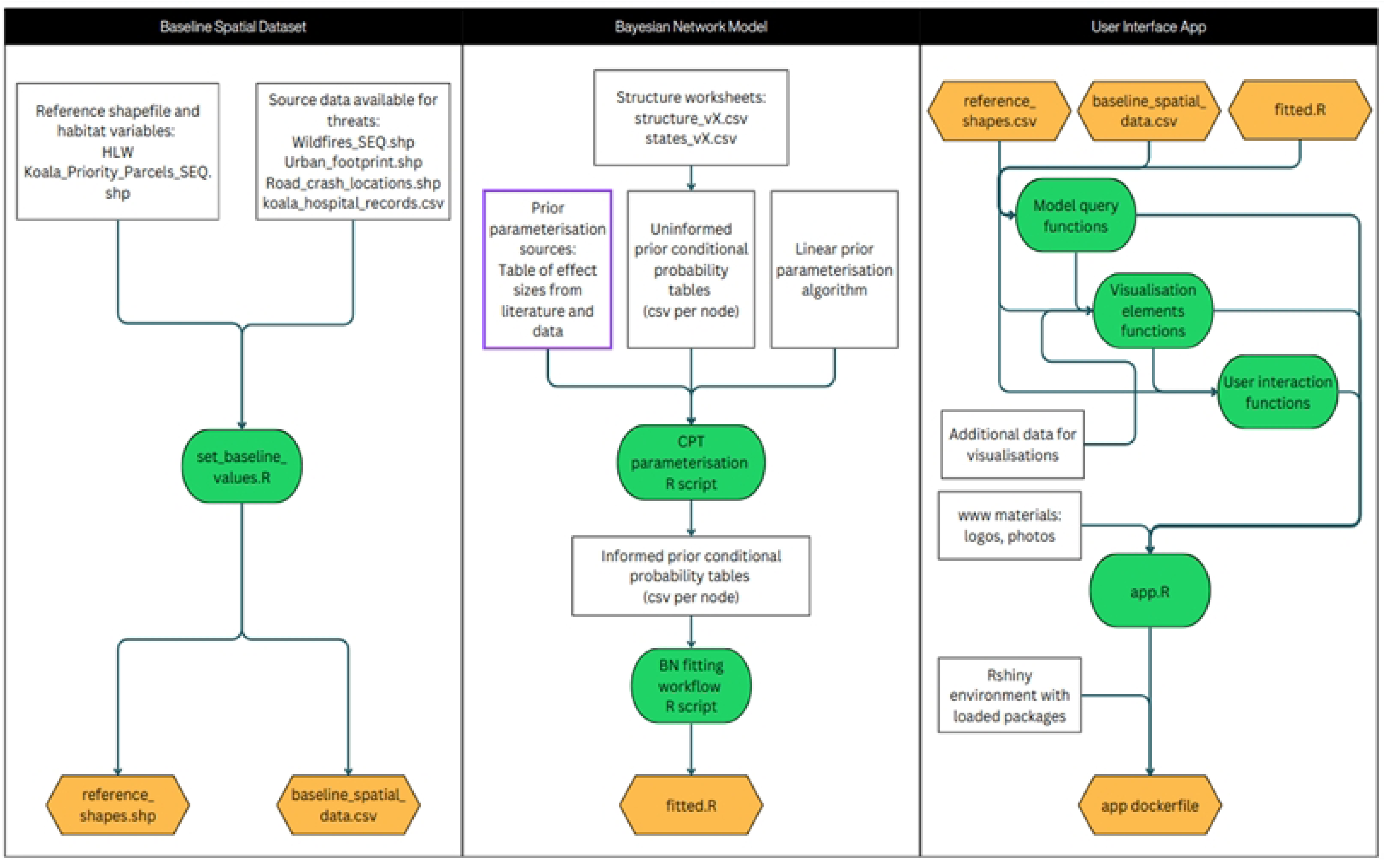
Components of the Koala Bayesian Network Platform (KBNP).

### KBNP results

Analysis of the geographical baseline data revealed variation in predicted KPP, habitat support, and threat level among local government areas (LGAs) in SEQ. LGAs in regional areas such as Somerset and the Scenic Rim had relatively high potential koalas. Urban LGAs with relatively small land parcels, such as Redland City, Logan City, and Noosa Shire, had smaller predicted potential koalas overall, as shown in Fig 4 (Left). Different LGAs had different levels of habitat support and threat level, as shown in Fig 4 (Right). Most LGAs had more land parcels with high habitat support than high threat levels, though some, like Redland City, had both high habitat support and high threats, while others, like South Burnett Regional, had low habitat support and low threats.

**Fig 4.**
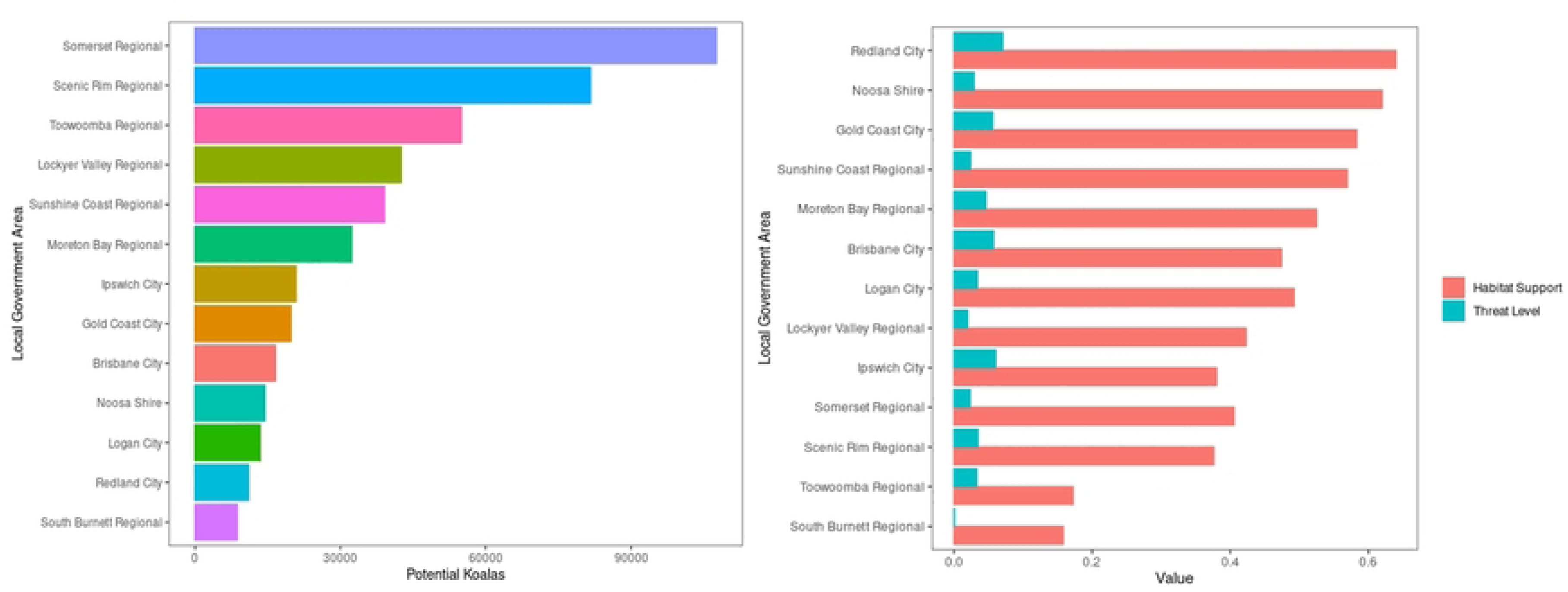
Predicted total potential koalas (left) and predicted habitat support and threat level per local government area (LGA).

LGAs also showed variation in baseline levels of different threats and habitat variables, as shown in Fig 5. Although baseline levels of threats fire, vehicle strike, and disease were relatively low in all LGAs, there was more variation in dog attack. Urban LGAs, including Brisbane City and Ipswich City, had relatively high levels of baseline dog attack risk. This finding may be partly explained by our baseline data for dog attack, which related to urban density. Baseline levels of habitat variables were more spread out across LGAs. Habitat connectivity was variable among LGAs, with Redland City showing the highest baseline value. Habitat size was low relative to other habitat variables, with larger tract sizes reported in Sunshine Coast, Somerset, Redland City, Noosa, and Gold Coast. Presence of koala tree species was high relative to other habitat variables, with more koala trees reported in Brisbane City, Logan City, Moreton Bay, Noosa, and Redland City.

**Fig 5.**
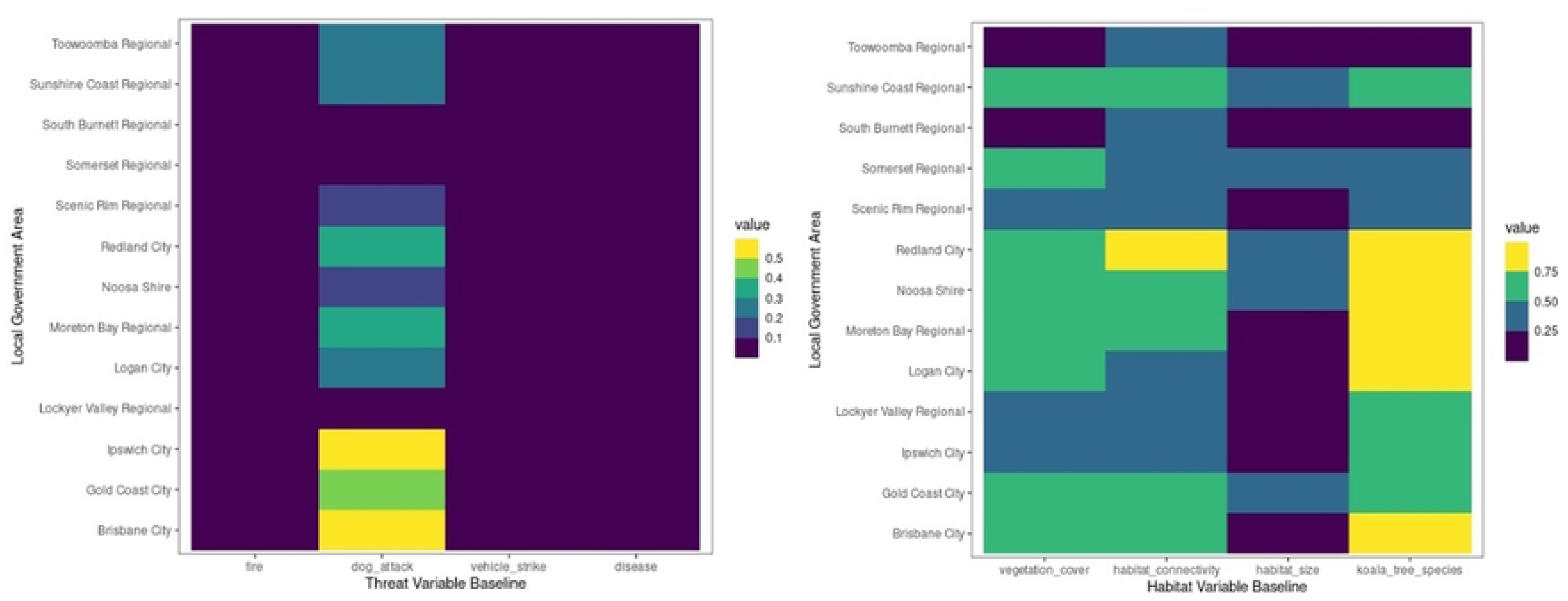
Relative levels of habitat and threat variables per local government area.

Maps of the habitat support and threat level in SEQ are shown in Fig 6. The uncertainty in habitat and threat predictions also varied among geographic regions. Running prediction queries on the BN 20 times per land parcel produced a distribution of habitat support and threat predictions per land parcel (Fig 6 Bottom). (To note, the 2019 wildfires influencing the high threat level observed in mapped landscape areas.)

**Fig 6.**
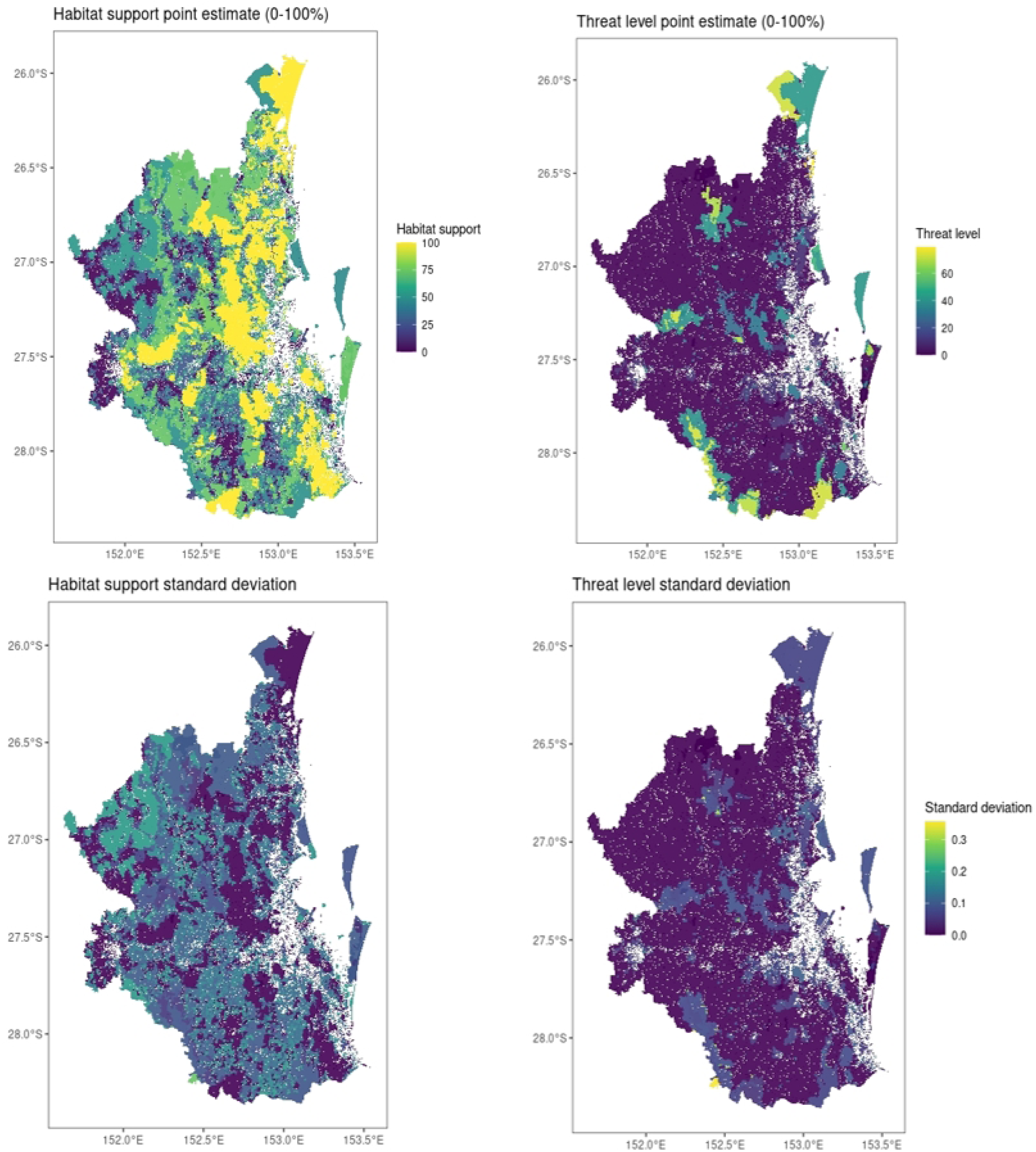
Maps of habitat support (Top left) and threat levels (Top right) and associated uncertainty (Bottom) per land parcel in SEQ.

The relative impact of the various indicators on habitat support and threat level is revealed in Fig 7. These marginal influence plots show how each variable individually contributes to the predicted probability of the target outcome, either high habitat support or high threat level, while holding other variables constant. The bars are ordered from most to least influential, and the colour indicates whether the influence is positive or negative. If the model is valid, we expect that the most influential variables would match the nodes on which the selected node is conditional in the network graph (Fig 2). As expected, habitat variables including ecology, vegetation cover, habitat size, and koala tree species had the highest influence on habitat support. Vehicle strike also appeared as a strong influencer in the habitat support model, likely because it frequently co-occurs in areas where koalas live. However, this does not imply a positive impact; rather, it reflects that regions with more habitat features also face higher exposure to road networks, which negatively affect koalas. The model predicted a current value of 46% habitat support across SEQ, with potential to reach 91%. As also expected, threat variables including vehicle strike, fire, disease, and dog attack had the highest influence on threat level, therefore negatively affecting koalas. The model predicted a current value of 4% threat level in SEQ; however, this low value should not be seen as evidence that threats are well-managed. It likely reflects the current probability of direct threat exposure and may underestimate long-term impacts, particularly in fragmented or developing areas. If key threats like vehicle strike, fire, and disease are not controlled, threat levels could rise to 76%.

**Fig 7.**
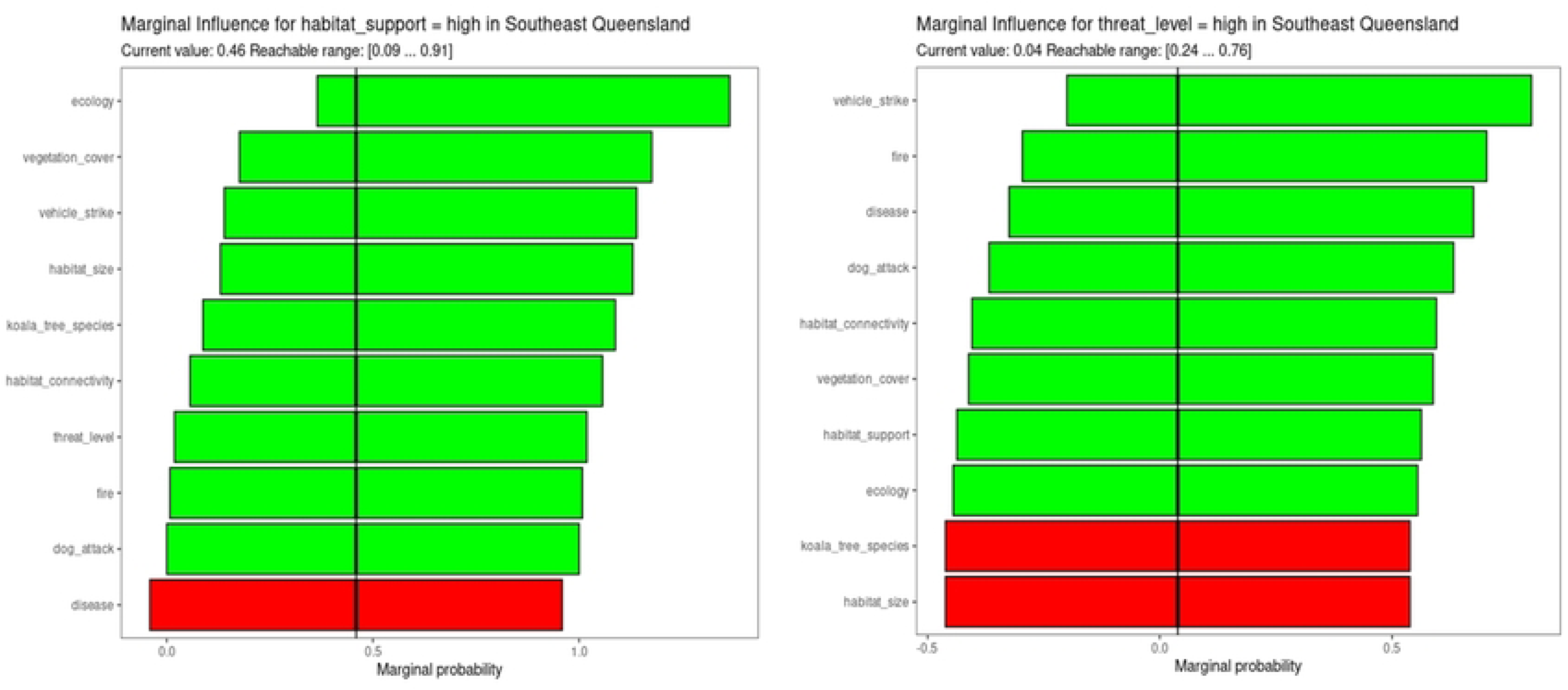
Marginal sensitivity for node habitat_support and reachable range (Left) and node threat_level and reachable range (Right) compared to baseline for SEQ.

Comparison of the habitat support predictions with the corresponding HLW DES scores for each land parcel in SEQ provided strong support for validity of the KBNP. Our habitat support score significantly predicted the HLW DES rating of habitat suitability for koalas, *R*^2^ = .803, *F* (1,121609) = 49950, *p* < .001 (Fig 8).

**Fig 8.**
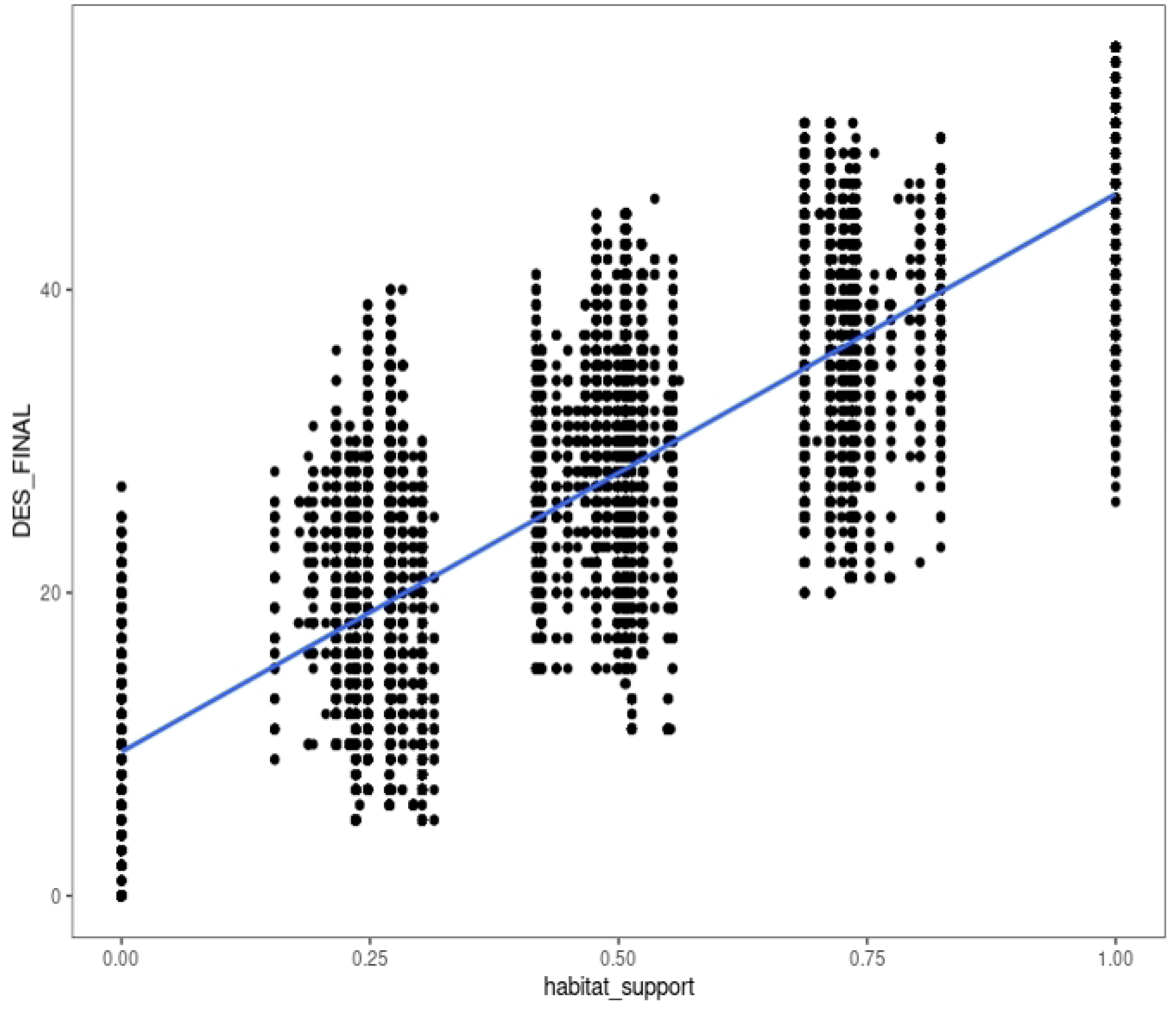
Predicted habitat support score (habitat_support) from the BN compared to HLW DES habitat suitability scores (DES_FINAL) for koalas.

## Discussion

This paper proposed a novel and integrative approach for supporting practical conservation management of koalas by diverse stakeholders including landowners, developers and government. The proposed KPP metric is based on the potential population of a species that could be supported in a specific geographic area. A complex systems approach is taken to develop the metric, taking into account the species and its required habitat characteristics, threats and potential conservation actions. A BN is employed to represent the system, thereby allowing for the ingestion of multiple data sources to probabilistically inform various components of the system and providing a location-specific output that is transformed to the desired metric. In addition to being shown to be reliable, explainable and responsive to local decisions around species threats and mitigating actions, this new measure provides a positive framing for managers, landholders and other stakeholders that can help to evaluate proactive decisions about conservation efforts under conditions of uncertainty and rapid change.

This research presents a strengths-based, collaborative, and future-oriented transdisciplinary metric and an accessible interactive digital platform that can aid in learning, identifying conservation strategies and support multi-way communication between stakeholders. One of the earliest wins from this work was the empowerment of decision makers to be able to visualize different scenarios in conservation efforts, and what the potential impacts are likely to be given a select set of actions. Local government and lobby groups found the platform particularly appealing as it provided opportunities for discussion and negotiation on alternatives especially around clearing of koala habitats for development.

The new KPP metric was demonstrated to effectively predict potential koala populations given habitat quality and threat levels in SEQ. The metric revealed substantial variation in the potential koalas supported throughout the region, and that different combinations of actions would be required to support viable koala populations in different regions. We conclude that management resources should be allocated in consultation with multiple stakeholders and considering local habitat and threat levels.

The KBNP tool has been developed in the context of koala conservation but is more widely applicable to other vulnerable species. It tackles the challenges of a consistent, standardised metric for discussing conservation actions, integrating disparate data sources and interactively evaluating the consequences of potential conservation actions at a local scale. It acknowledges known difficulties associated with documenting actual koala numbers in the wild, rapid habitat conversion, long-lag time for population recovery, unknowns about the species adaptation to changes, and the epidemiology of disease and impacts on target populations, all of which make it challenging to demonstrate the direct and indirect impacts of the different efforts undertaken for conservation. These challenges are further compounded by the reliance on baseline data, which requires regular updating to ensure the accuracy and relevance of conservation assessments. Although the system is designed to incorporate new data, it is essential to recognise the inherent limitations and variability in data quality. The tool directly addresses a need to provide a way to showcase incremental gains/benefits as a result of stakeholders’ positive actions for conservation, even if they can’t actually document an increase in the number of species actually observed.

Key conservation decisions must balance benefit, cost and impact. Potential actions are typically constrained by funding, feasibility and timeliness, as well as other priorities for funding [22]. In a resource-constrained environment, an optimal combination of actions is one with maximum impact is achieved given the limited resources.

The choice of which action or set of actions is “optimal” can be different for different stakeholders. Responsibilities and costs of each solution will be distributed unevenly among government levels, private individuals, and conservation groups [21]. Conservationists must therefore balance diverse public and private interests [31], and cooperation among diverse stakeholders is key to identifying an appropriate mix of management strategies [23]. The systems tool, the associated population potential metric and the interactive digital platform presented in this paper provides an appealing approach to addressing these pressing needs.

Future work could focus on key areas to enhance the model’s applicability. This includes the transfer of the model to other regions to test its scalability, and broader validation through comparisons with other KPP metrics. An extension to Dynamic Bayesian Networks (DBN) could enable “what-if” scenario analyses over time, assessing the iterative impacts of management actions. Finally, as with any statistical model employed in practice ongoing stakeholder user testing across diverse groups is necessary to refine the model’s usability and ensure it meets the needs of conservation efforts.

## Acknowledgments

The authors acknowledge the following contributions.

Melissa Walker, HLW, and Michael Petter, formerly of SEQ Catchments, for their support in spatial and mapping decision-making, including essential habitat mapping for species.

The Koala Habitat Partnerships and Programs Team of the Department of Environment, Science, and Innovation for their interest in the work and their support of the project.

More broadly, the authors acknowledge citizen scientists of SEQ for monitoring koalas and lobbying for greater koala conservation for over 20 years, as well as the First Nations People of SEQ who are actively caring for country and working on Koala conservation.

